# Turtle body size evolution is determined by lineage-specific specializations rather than global trends

**DOI:** 10.1101/2022.12.13.520223

**Authors:** Bruna M. Farina, Pedro L. Godoy, Roger B.J. Benson, Max C. Langer, Gabriel S. Ferreira

## Abstract

Organisms display considerable variety of body sizes and shapes, and macroevolutionary investigations help to understand the evolutionary dynamics behind to such variations. Turtles (Testudinata) show great body size disparity, especially when their rich fossil record is accounted for. We explored body size evolution in turtles, testing which factors might influence the observed patterns and evaluating the existence of long-term directional trends. We constructed the most comprehensive body size dataset for the group to date, tested for correlation with paleotemperature, estimated ancestral body sizes, and performed macroevolutionary model-fitting analyses. We found no evidence for directional body size evolution, even when using very flexible models, thereby rejecting the occurrence of Cope’s rule. We also found no significant effect of paleotemperature on overall through-time body size patterns. In contrast, we found a significant influence of habitat preference on turtle body size. Freshwater turtles display a rather homogenous body size distribution through time. In contrast, terrestrial and marine turtles show more pronounced variation, with terrestrial forms being restricted to larger body sizes, up to the origin of testudinids in the Cenozoic, and marine turtles undergoing a reduction in body size disparity after the extinctions of many groups in the mid-Cenozoic. Our results therefore suggest that long-term, generalized patterns are probably explained by factors specific to certain groups and related at least partly to habitat use.

## 1 INTRODUCTION

Organisms have evolved a remarkable disparity of body plans, sizes, and functions (Smith et al., 2016), and the relationship between phenotypic evolution and species diversification is a widely discussed subject in evolutionary biology (e.g., Cooney & Thomas, 2021; Stanley, 1973). Body size in particular has been shown to affect several traits in organisms, especially life history (Brown et al., 1993; White et al., 2022), metabolic rates (Blanckenhorn, 2000; Clauss et al., 2007; D’Amico et al., 2001), and ecology (Brown & Maurer, 1986; Smith et al., 2018; White et al., 2007). For such reasons, the topic has always intrigued researchers and its role in microevolution (i.e., natural and artificial selection) and macroevolution (e.g., acquisition of new traits or production of ecological opportunities) has been extensively debated (Blanckenhorn, 2000; Maurer et al., 1992; Peters, 1983; Schmidt-Nielsen & Knut, 1984; Stanley, 1973). The evolution of body size is commonly explained in the light of different hypotheses that attempt to elucidate patterns of disparity, such as Cope’s rule (or Deperet’s rule; Stanley, 1973), a hypothesized tendency lineages to evolve towards larger body sizes (Cope, 1896). Although directional trends of increasing body size have been identified in a few groups, including medium to large-bodied mammals (Alroy, 1998) and pterosaurs (Benson et al., 2014), non-directional patterns are present in many other groups (Benson et al., 2018; Godoy et al., 2019; Laurin, 2004; Moen, 2006). The generalization of hypothesized evolution towards large body sizes has also been questioned, because there is no apparent need for selection to produce ever larger sizes in most lineages (Gould, 1988).

In spite of the presence of hypotheses to explain evolutionary variation of body size in ectothermic vertebrates, few groups have been studied in a comprehensive manner (but see Gearty & Payne, 2020; Godoy et al., 2019; Heim et al., 2015; Smith et al., 2016), with most studies focusing on endothermic groups such as mammals and dinosaurs, including birds (Benson et al., 2014, 2018; Cooper & Purvis, 2010; Cullen et al., 2020; Gearty et al., 2018; Kubo et al., 2019; Raia & Meiri, 2011). Turtles, in particular, have a rich fossil record and relatively stable phylogenetic relations, which provide a reliable framework for macroevolutionary studies. The presence of fossils is particularly important because the inclusion of deep-time data can have a large effect on the outcomes of macroevolutionary analyses.

Turtles include 357 living species (Rhodin et al., 2021), but the fossil record of crown-group Testudines, and that of the more inclusive group Testudinata, reveals a much richer history (Gaffney et al., 2006; W. G. Joyce et al., 2013). The early evolution of the turtle stem-lineage is thought to have occurred in terrestrial habitats (Joyce, 2017; Joyce & Gauthier, 2004; Lautenschlager et al., 2018), but aquatic habits evolved towards the crown-group, and represents the ancestral condition for Testudines (Joyce, 2017; Joyce & Gauthier, 2004). Therefore, possibly reflecting the different habitats occupied through time (although not only for that reason), the group shows considerable morphological disparity in their limbs, skull, carapace shape, and body size (Benson et al., 2011; Dickson & Pierce, 2019; Foth et al., 2017; Hermanson et al., 2022; Jaffe et al., 2011; Joyce & Gauthier, 2004; Lautenschlager et al., 2018; Vlachos & Rabi, 2018).

Considering the relatively low extant diversity of turtles, particularly when compared to mammals, birds, or squamates, the body size disparity of the group is striking. The smallest living testudine, *Homopus signatus*, has an adult carapace length of about 100 mm, and the largest one, *Dermochelys coriacea*, reaches more than 2200 mm (Rhodin et al., 2021). Furthermore, fossils display an even broader range of body sizes, including the South American *Stupendemys geographicus*, with a carapace length of more than 2800 mm (Cadena et al., 2020). This emphasizes the importance of including the available fossil diversity when characterizing patterns of body size evolution in Testudinata (and see Finarelli & Flynn, 2006; Fritz et al., 2013), which has been largely disregarded in most previous attempts (Eastman et al., 2011; Jaffe et al., 2011; Moen, 2006; Uyeda & Harmon, 2014).

Using data from living species only, previous studies presented several hypotheses to explain the observed body size variation of turtles. For example, it has been suggested that such variation is intrinsically related to habitat, with marine species and island tortoises usually possessing larger sizes than freshwater and mainland taxa (Jaffe et al., 2011). This resembles the large body size attained by marine mammals (e.g. Gearty et al. 2018) and, in some aspects, the ‘island rule’ seen in some mammals, reptiles, and birds (Lomolino, 2005). However, Uyeda & Harmon (2014) analyzed turtle body size using unconstrained evolutionary models and suggested that the scenario for turtle optimal body size is more complex than simple differences in habitats, with multiple macroevolutionary body size shifts along the tree. Furthermore, analysis of tortoise (Testudinidae) body size, including fossils, did not find support for the island effect (Vlachos & Rabi, 2018). Moen (2006) tested for evolutionary trends in extant cryptodires, but found no support for directional body size evolution. Moreover, it is not currently clear how temperature influenced long-term body size patterns in turtles, with ongoing discussion about the group following overall rules such as a latitudinal gradient, with larger body sizes seen in colder regions (Angielczyk et al., 2015; Ashton & Feldman, 2003). Yet, except for Vlachos & Rabi (2014), these hypotheses have yet to be tested in a framework including both extinct and extant taxa. In the present study, we compiled the largest body size dataset ever assembled for Testudinata, which was used to investigate the tempo and mode of body size evolution in the group, as well as test for possible biotic and abiotic drivers.

## 2. METHODS

### 2.1. Body size data

Straight-line maximum dorsal carapace length (SCL) was used as a proxy for turtle body size (Jaffe et al., 2011). Aiming to maximize sampling, we also used linear regressions to estimate SCL from the ventral skull length (measured from the rostral tip of the premaxillae to the caudal tip of the occipital condyle) for some specimens lacking carapace. About 7.5% of the SCL data in our dataset was estimated from the ventral skull length. Measurements were collected from photographs (personal archive or the literature), using software ImageJ (Schneider et al., 2012). The final dataset includes body size data for 795 taxa, considerably more than in previous studies (e.g., Jaffe et al., 2011 = 226 taxa; Angielczyk et al., 2015 = 245 taxa; Vlachos & Rabi, 2018 = 59 taxa; Moen, 2006 = 201 taxa). Additionally, we also collected habitat preference and chronostratigraphic information for these same taxa using the literature and the Paleobiology Database (PBDB).

### 2.2. Supertree construction and time-calibration

To account for major uncertainties within the phylogenetic relations of the main groups of Testudinata, two informal supertrees were manually assembled using Mesquite version 3.61 (Maddison & Maddison, 2018). These were based on two phylogenetic hypotheses, Evers et al. (2019) and Sterli et al. (2018), hereafter referred to as “Ev19” and “St18”, respectively. The most significant differences between the two supertrees are the positions of Protostegidae and Thalassochelyidia (sensu Joyce et al., 2021). Protostegids are stem-Chelonioidea and Thalassochelyidia are stem-Pleurodira in “Ev19”, whereas both groups belong to the turtle stem-lineage in “St18” (in which they are originally represented only by *Santanachelys gaffneyi* and *Solnhofia parsoni*, respectively). Less inclusive groups were positioned based on several additional hypotheses (Table A1). Each supertree includes a total of 846 taxa, 659 of which are shared with our body size dataset.

Both supertrees were time-scaled using Bayesian inference under a fossilized birth death (FBD) process (Heath et al., 2014; Stadler, 2010), performed with MrBayes version 3.2.7 (Ronquist et al., 2012). We used R (version 4.0.2; R Core Team, 2021) package *paleotree* (Bapst, 2012) to create a MrBayes command for time-calibration analyses. The function *createMrBayesTipDatingNexus()* allows the use of “empty” morphological matrices in clock-less tip-dating analyses (Bapst, 2012). The two supertrees (“Ev19” and “St18”) were entered as topological constraints (i.e., for two separate time-scaling analyses) and data on occurrence times (=tip ages) were obtained from the primary literature and supplemented by the PBDB. Four taxa formed the outgroup (*Eunotosaurus africanus*, *Eorhynchochelys sinensis, Pappochelys rosinae*, and *Odontochelys semitestacea*) and, based on their age (as well as that of the oldest turtles), the root of the tree was defined between the Kungurian and Roadian stages of the Permian (283.5 and 268.8 million years ago, Ma) using a uniform distribution, as this would represent a maximum possible age for the origin of the group. The polytomies were randomly resolved. Two runs of MCMC analyses, with four chains each, were set for 20,000,000 generations, with 25% burn-in. Convergence of both runs were verified when values of potential scale reduction factors approached 1.0 and average standard deviation of split frequencies was below 0.01. For both supertrees, we used either the maximum clade credibility (MCC) tree or a set of 10 randomly selected trees to perform subsequent analyses.

### 2.3. Characterizing body size patterns in Testudinata

The entire body size dataset of 795 taxa was used to construct body size through-time plots. Welch’s Two Sample t-tests (Welch, 1947) were used to assess significant changes across different time intervals (i.e., Triassic, Jurassic, Early Cretaceous, Late Cretaceous, Paleogene, Neogene, and Quaternary), focusing on mean body size and disparity, using the standard deviation as a metric of body size disparity. To assess the influence of ecology on the body size distribution, habitat preference information (i.e., terrestrial, freshwater, and marine) was also incorporated into the body size through-time plots. To further test the influence of ecology, we used analysis of variance (ANOVA), performed with R function *aov()*, as well as the RRPP approach (randomizing residuals in a permutation procedure; Adams & Collyer, 2018), which accounts for phylogenetic dependency (i.e., “phylogenetic ANOVA”), performed with the *lm.rrpp()* function and using the MCC tree of each supertree (“Ev19” and “St18”).

We also tested for the presence of phylogenetic signal in the body size data using the R function *phyloSignal()* (Keck et al., 2016), using 10 randomly selected trees from the posterior distribution of trees of both supertrees. We used 1,000 replicates and estimated Pagel’s lambda (λ) as our metric of phylogenetic signal given that this index is robust when using trees with poorly resolved branch length information (Molina-Venegas & Rodríguez, 2017; Münkemüller et al., 2012).

To further characterize body size evolution within Testudinata, we used maximum likelihood to estimate ancestral body sizes under Brownian motion, using the *fastAnc()* function of the R package *phytools* (Revell, 2012). Ancestral body size reconstructions were performed with both the complete supertrees (i.e., using the MCCT trees with all 659 taxa, including fossils and extant species) and a subtree with only extant taxa (i.e., dataset reduced to 312 taxa).

### 2.4. Testing for the presence of Cope’s rule

To test if Cope’s rule played an important role in turtle body size evolution, we fitted different evolutionary models to our body size data in both supertrees. To account for temporal and phylogenetic uncertainties, 10 time-scaled versions of each alternative supertree (“Ev19” and “St18”) were used.

We fitted four uniform phenotypic models to our data, starting with the uniform Brownian Motion (BM) model, in which body size undergoes an unconstrained, single-rate random walk along phylogenetic lineages, resulting in diffusive evolutionary expansion (Felsenstein, 1973, 1985; Freckleton & Harvey, 2006). This pattern is consistent with several possible causes, including genetic drift or wandering adaptive optima (Felsenstein, 2003), between which genetic drift seems a less likely explanation at macroevolutionary scales. The model has two parameters: sigma squared (σ ), which indicates evolutionary rate, and the root state of the trait at time zero, sometimes represented by *X(0)* (Felsenstein, 1973)

We also fitted three other uniform models: (1) the “trend” model, which is a modification of the BM model that incorporates a parameter (µ ) describing an uniform directional trend along all branches of the phylogeny (Pagel, 2002); (2) the EB model (also known as “ACDC model”; Blomberg et al., 2003), in which lineages experience a burst of rapid increase in trait variation in the beginning of their evolutionary history, followed by a deceleration (Harmon et al., 2010); (3) the Ornstein-Uhlenbeck (OU) model, which incorporates attraction of trait values (represented by the α parameter) towards an optimum (θ) (Butler & King, 2004; Hansen, 1997). In the case of the OU model fitted here, the parameters (α and θ) were not allowed to vary along the tree.

We also fitted 13 non-uniform trend-like models to our data. Unlike uniform “trend” model, these multi-regime models allow the μ parameter – the amount of directional change in a trait through time (Hunt & Carrano, 2010; Pagel, 2002) – to vary along the tree in temporal or node shifts (“time-shift model” and “node-shift model”, respectively). We fitted “time-shift” models (which allow shifts in all branches after a determined point in time) allowing the number of shifts to vary from one to three; and “node-shift” models (which allow shifts in some branches) allowing the number of shifts to vary from one to ten, resulting in a total of 13 multi-trend models (three “time-shift” and ten “node-shift” models). In the present study, both uniform trend and multi-trend models were fitted as a representation of the Cope’s rule, given that it is described as a multi-lineage directional trend towards larger sizes (Cope, 1887, 1896; Stanley, 1973). Moreover, among mammals, the foundational example of Cope’s rule, directional body size evolution is only present in some lineages (e.g., Alroy, 1999), more consistent with a “node-shift” model, with multiple independent origins of directional evolution.

Akaike’s information criterion for finite sample sizes (AICc) was used for the selection of the best fit (Akaike, 1974). Model-fitting analyses were performed using R package *geiger* (Harmon et al., 2008) and the scripts made available by Benson et al. (2018) for fitting the multi-trend models.

### 2.5. Influence of paleotemperature

We used regressions to test for the possible influence of global paleotemperature on turtle body size (795 taxa). As a proxy for paleotemperature, we compiled δ^18^O data (lower δ^18^O values indicate higher environmental temperature) from two different sources. Firstly, we used isotopic data collected in tropical regions by Prokoph et al. (2008), who assembled isotopic information from marine organisms, extending from precambrian to recent. Further, we also used data from Zachos et al. (2008), which compiled information about isotopic ratios in foraminifer shells from the Maastrichtian to the recent. We tested for correlation between both temperature curves and our body size indices, including maximum, minimum, and mean body size, as well as body size disparity (= standard deviation).

Correlation between body size data and paleotemperature was initially assessed using ordinary least squares (OLS). Additionally, to avoid potential issues created by temporal autocorrelation, we used generalized least squares with a first-order autoregressive model incorporated (tsGLS; Fox & Weisberg, 2018), using the R package *nlme* (Pinheiro et al., 2022). The data was divided into time intervals, using approximately equal-length (∼9 million years) stratigraphic time bins (from Mannion et al., 2015). For each time bin, we calculated body size indices (disparity, maximum, minimum, and mean body size) and weighted mean δ^18^O values.

## 3. RESULTS

### 3.1. Ancestral state reconstructions

Ancestral state reconstructions based on both supertrees (“Ev19” or “St18”; Figures A3 and 1) show similar results. For this reason, the description below is based solely on “St18” (Figure 1).

**Figure 1.**
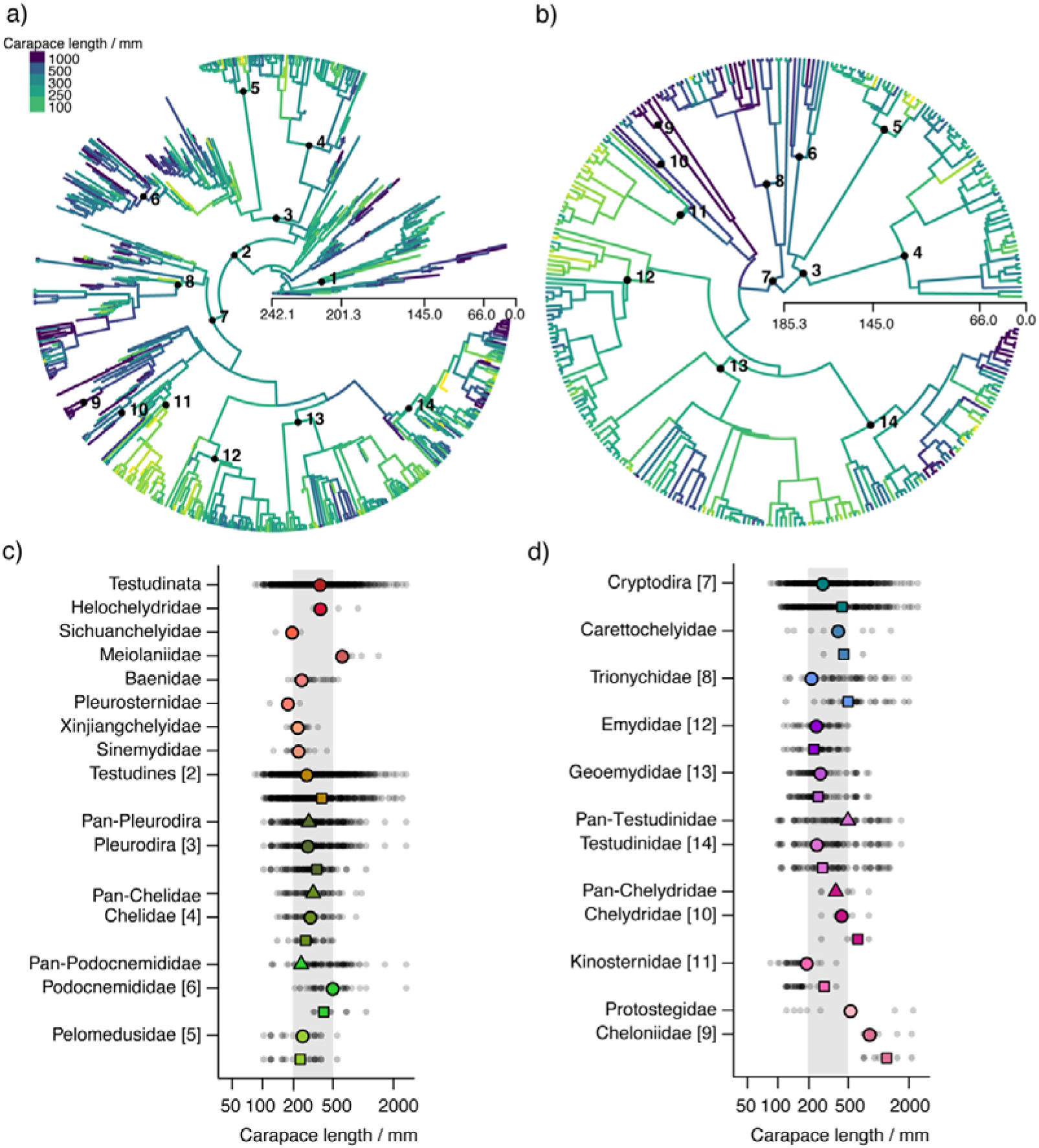
Ancestral body sizes (log_10_ maximum dorsal carapace length in millimeters) mapped onto Testudinata phylogenies, with (a) complete tree and (b) extant-only subtree. Ancestral body size for different taxonomic groups (c and d); small grey dots indicate all taxa within that lineage; colored triangles represent the ancestral estimates of stem-groups; circles represent ancestral estimates of the crown-groups and squares represent ancestral estimates of the crown-groups without the fossil taxa. Grey area indicates sizes between 200 and 500 mm. The numbers in the panels represent groups: 1. Paracryptodira; 2. Testudines; 3. Pleurodira; 4. Chelidae; 5. Pelomedusidae; 6. Podocnemididae; 7. Cryptodira; 8. Trionychidae; 9. Cheloniidae; 10. Chelydridae; 11. Kinosternidae; 12. Emydidae; 13. Geoemydidae; 14. Testudinidae.

**Table 3.** Results of ANOVA and PhyloANOVA exploring the relationship between Testudinata body size (log_10_ maximum dorsal carapace length in millimeters) and different ecological categories: terrestrial (n=145), freshwater (n=563), and marine (n=87). Pairwise comparisons between ecological categories also shown. Abbreviations: Df, degrees of freedom; SS, sum of squares; MS, mean squares. *Significant at alpha = 0.05.

| ANOVA results | Df | SS | MS | F-value | p-value |
| --- | --- | --- | --- | --- | --- |
| **ANOVA** | 2 | 4.77 | 2.3868 | 36.55 | <0.0001* |
| **PhyloANOVA** | 2 | 0.0323 | 0.0161295 | 1.8106 | 0.109 |
| Pairwise comparison (p-values) | Marine-Fresh. |  | Terrestrial-Fresh. |  | Terrestrial-Marine |
| **ANOVA** | <0.0001* |  | <0.0001* |  | 0.502 |
| **PhyloANOVA** | 0.801 |  | 0.04* |  | 0.176 |

When fossil taxa are included in the reconstructions, ancestral size estimates for most major (= more diverse) turtle subgroups (e.g., Testudines, Pleurodira, Chelidae, Pelomedusidae, Cryptodira, Trionychidae, and Testudinidae) were broadly similar to one another, with SCL values between 500 and 230 mm (Figure 1c and d). Similar ancestral values are also seen among stem-turtles, with paracryptodires remaining within approximately this body size range throughout their evolution and meiolaniids increasing their body size over time, from an ancestral body size estimated in 645 mm (Figure 1). Among pleurodires, several extinct branches splitting before the origin of Podocnemididae show smaller body sizes (between 100 and 250 mm; Figure 1a and c), although the estimated ancestral body size for Podocnemididae is larger (between 300 and 500 mm; Figure 1c). Within Cryptodira, crown-groups Testudinidae, Geoemydidae, and Emydidae have similar ancestral body sizes (about 250 mm; Figure 1d). Chelonioidae show ancestral body sizes above 500 mm, and Kinosternidae was one of the few main clades with an estimated ancestral body size close to (or slightly above) 200 mm.

The inclusion of fossils affects ancestral body size reconstructions for most major lineages (compare square symbols [ignoring fossils] to circles [including fossils] in Figure 1c, d). No specific directional influence is noted when paleontological data is included. For some groups (e.g., Chelidae, Podocnemididae, Emydidae, Geoemydidae), the inclusion of extinct taxa results in a slight increase in estimated body sizes, whereas for others (e.g., Pelomedusidae, Trionychidae, Testudinidae, Chelydridae, Kinosternidae, Cheloniidae), a decrease is observed. The magnitude of this effect varies; the largest changes were seen in the nodes circumscribing Cryptodira, Trionychidae, Chelydridae, Kinosternidae, and Cheloniidae (Figure 1). It is worth noting that even a slight increase in fossil sampling changed the estimates in relation to previous studies. For instance, the ancestral body size for Pan-Testudinidae – based on 78 taxa (53 living and 25 extinct) – is 496.3 mm. Larger than the 370 mm estimated by Vlachos & Rabi (2018), which included 59 taxa (23 living and 36 extinct).

### 3.2. Model fitting

The AICc scores for all the evolutionary models fitted to the turtle trees and body size data show an overwhelmingly stronger support (i.e., lower AICc values) for the uniform OU model, even when compared to the non-uniform multi-trend models. Consistently, this stronger support for the OU model was found when using both “Ev19” and “St18” topologies (Table 1). These results rule out the presence of trend-like processes (either uniform or multi-trend) in the body size evolution of Testudinata, at least when the entire tree is considered.

**Table 1.** Results of model-fitting analyses, depicting model parameters and AICc scores for the models fitted to our body size dataset of Testudinata (log_10_ maximum dorsal carapace length) and 10 time-calibrated trees for each of the two initial supertree topologies (“St18”, based on the hypothesis of Sterli et al. [2018], and “Ev19”, based on the hypothesis of Evers et al. [2019]). Models: **BM** (Brownian Motion model), **EB** (Early Burst/ACDC model), **OU** (Ornstein-Uhlenbeck model) and **“best” trend** (the model with best fit [AICc scores] among the 14 trend-like models fitted [1 uniform and 13 non-uniform models], which in the case of “St18” is represented by the non-uniform trend model with 4 “time-shifts”, and in the case of “Ev19” is represented by the uniform trend model). Mean values of model parameters are shown for the 10 time-calibrated trees: σ**^2^** (sigma squared, the Brownian variance or rate parameter), X***(0)*** (estimated trait value at the root of the tree, also known as Z_0_), and μ (the trend parameters, describing a uniform directional trend along all branches of the phylogeny, with the number of parameters varying according to the number of shifts). The mean **AICc** scores indicate overwhelming support (i.e., lower AICc values) to the OU model over the other models.

| Supertree | St18 |  |  |  | Ev19 |  |  |  |
| --- | --- | --- | --- | --- | --- | --- | --- | --- |
| Parameter | BM | EB | OU | “best” trend | BM | EB | OU | “best” trend |
| **AICc** | 782.09 | 784.11 | 96.69 | 665.15 | 976.62 | 978.64 | 105.28 | 861.80 |
| $\sigma^2$ | 0.0142 | 0.0142 | 0.5833 | 0.0128 | 0.0185 | 0.0185 | 1.7895 | 0.432 |
| $X(0)$ | 2.6764 | 2.6764 | 2.5124 | 2.6148 | 2.669 | 2.669 | 2.5113 | 2.6038 |
| $\mu_1$ | -- | -- | -- | 0.0023 | -- | -- | -- | 0.0024 |
| $\mu_2$ | -- | -- | -- | -0.7601 | -- | -- | -- | -- |
| $\mu_3$ | -- | -- | -- | -0.0238 | -- | -- | -- | -- |
| $\mu_4$ | -- | -- | -- | -0.3237 | -- | -- | -- | -- |
| $\mu_5$ | -- | -- | -- | -0.7850 | -- | -- | -- | -- |

### 3.3. Correlation with paleotemperature

No significant correlation was observed between global paleotemperature and mean, minimum, or maximum body size of Testudinata through time (Table A2). However, we did find a significant, but weak correlation between body size disparity (= standard deviation of body sizes) and paleotemperature (Table 2; Figure 2), with disparity increasing at lower temperatures (i.e., higher δ^18^O values). Although significant, this correlation is relatively weak when palaeotemperature data from Prokoph et al. (2008) is used, but it becomes slightly stronger when using data from Zachos et al. (2008), which is restricted to the Late Cretaceous-Recent time interval. This may suggest that the influence of environmental temperatures on turtle body size was stronger during the Cenozoic.

**Table 2.**
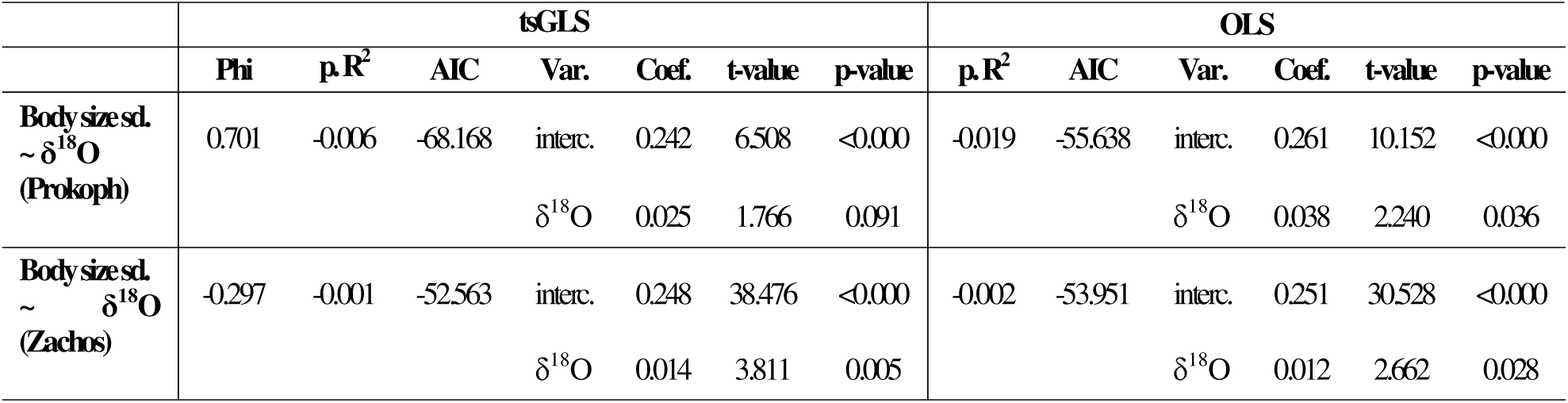
Results of time series generalized least squares (tsGLS) and ordinary least squares (OLS) regressions using turtle body size disparity and paleotemperature data. Body size disparity (standard deviation of log_10_ maximum dorsal carapace length in millimeters) and paleotemperature (δ^18^O data as a proxy for paleotemperature from two different sources: Prokoph et al. [2008] and Zachos et al. [2008]) data was divided into time bins (24 time-bins when the Prokoph et al. [2008] δ^18^O data was used and 10 time-bins when the Zachos et al. [2008] data was used). Abbreviation: **sd.**, standard deviation; **p.R^2^**, Nagelkerke pseudo R-squared; **Var.**, variable; **interc.**, intercept; **Coef.**, coefficient.

|  | tsGLS |  |  |  |  |  |  | OLS |  |  |  |  |  |
| --- | --- | --- | --- | --- | --- | --- | --- | --- | --- | --- | --- | --- | --- |
|  | Phi | p.R<sup>2</sup> | AIC | Var. | Coef. | t-value | p-value | p.R<sup>2</sup> | AIC | Var. | Coef. | t-value | p-value |
| **Body size sd.**<br>**~ $\delta^{18}\text{O}$**<br>**(Prokoph)** | 0.701 | -0.006 | -68.168 | interc. | 0.242 | 6.508 | <0.000 | -0.019 | -55.638 | interc. | 0.261 | 10.152 | <0.000 |
| | | | | $\delta^{18}\text{O}$ | 0.025 | 1.766 | 0.091 | | | $\delta^{18}\text{O}$ | 0.038 | 2.240 | 0.036 |
| **Body size sd.**<br>**~ $\delta^{18}\text{O}$**<br>**(Zachos)** | -0.297 | -0.001 | -52.563 | interc. | 0.248 | 38.476 | <0.000 | -0.002 | -53.951 | interc. | 0.251 | 30.528 | <0.000 |
| | | | | $\delta^{18}\text{O}$ | 0.014 | 3.811 | 0.005 | | | $\delta^{18}\text{O}$ | 0.012 | 2.662 | 0.028 |

**Figure 2.**
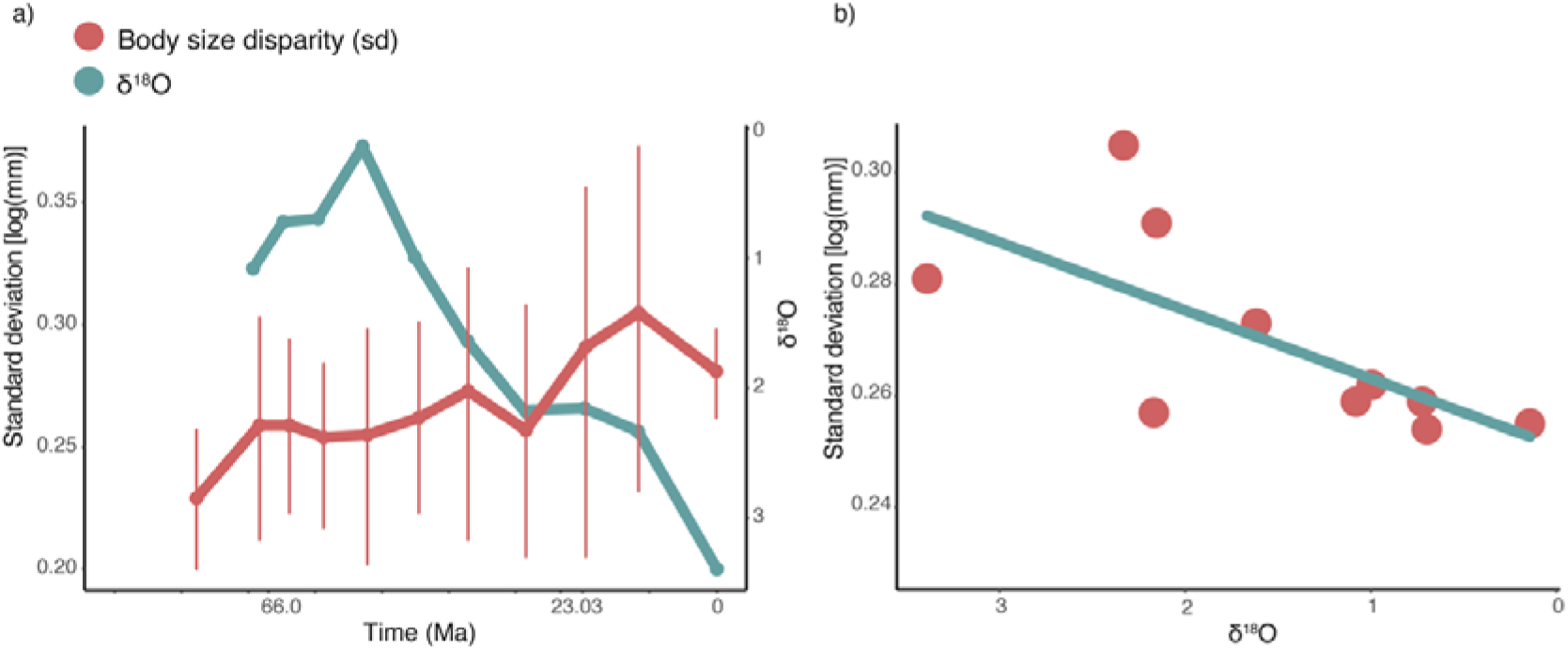
(a) Through-time patterns of Testudinata body size disparity (standard deviation of log_10_ maximum dorsal carapace length in millimeters) and paleotemperature (δ^18^O isotopic data from Zachos et al. [2008]) during the last ∼70 Ma. Error bars were calculated by bootstrapping the disparity data 500 times. δ^18^O is used as proxy for paleotemperature and is inversely proportional to temperature. (b) Linear regression (OLS) between turtle body size disparity and δ^18^O data (regression results shown in Table 2).

### 3.4. Differences among ecological habitats

Freshwater is the most common habitat occupied by turtles from the Jurassic onwards, and this category is therefore the most influential to the aggregated pattern of turtle body size variation through time. No significant changes in either disparity or mean body size of freshwater turtles occurred since the Late Cretaceous (Figure 3b, d). A significant increase in mean body size in freshwater turtles was identified between the Early (234 mm) and the Late Cretaceous (346 mm) (p-value = 0.001146). In general, freshwater turtles are more frequently represented among the smallest body sizes, and only rarely among the largest ones (Figure 3b).

**Figure 3.**
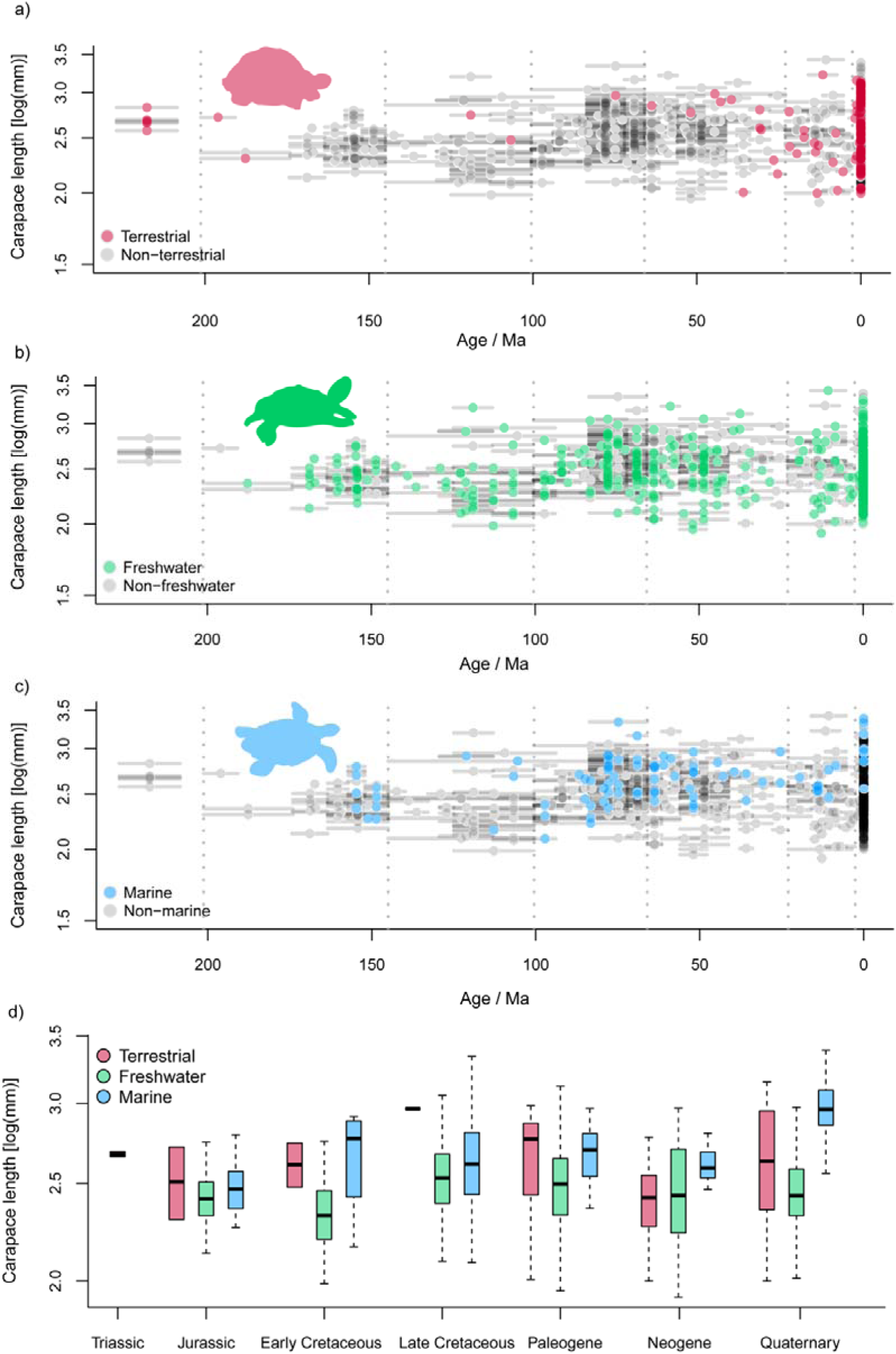
Temporal distribution of body sizes (log_10_ maximum dorsal carapace length in millimeters) in turtles for different ecological habitats. Gray dots represent all taxa, whereas colored dots represent taxa subdivided into three ecological categories. a) terrestrial taxa (red dots); b) freshwater taxa (light green dots); c) and marine taxa (light blue dots). Horizontal grey segments represent the range of occurrence of each taxon; d) Boxplot showing body size of different ecological categories divided into time intervals. Silhouettes adapted from Jaffe et al. (2011). We found that ecological habitat is significantly linked to body size in turtles using ANOVA (Table 3). However, when the phylogenetic structure of the data is taken into account (using phylogenetic ANOVA; Table 3), this association is only significant for the freshwater and terrestrial ecological categories. Moreover, for all trees tested here, we found a strong phylogenetic signal in body size data (λ > 0.856; P <0.001).

Diversity in terrestrial turtles is low until the Paleogene, when the first tortoises (Testudinidae) appear in the fossil record (Figure 3a). The mean body size of terrestrial turtles is consistently larger than that of freshwater species through time, except during the Neogene (Figure 3d). From the Triassic to the Late Cretaceous, terrestrial turtles experienced a significant increase in mean body size (from 272 to 443 mm; p-value = 0.02865). However, body size disparity is low, as expected from the low number of species, and the group is mostly represented by medium or large-bodied forms (Figure 3a).

The first marine turtles appeared at the end of the Jurassic, with body sizes like those of other groups (Figure 3c). There is a noteworthy (although not significant; p-value = 0.3872) increase in the mean body size of marine turtles from the Jurassic to the Early Cretaceous, and a small drop in the Late Cretaceous (Figure 3 c-d), but the latter interval witnessed the highest variability in body sizes among marine turtles in the series (Figure 3c). After the K-Pg transition, the range of body sizes decreased substantially, although the mean remains similar. After that, from the Neogene to the Quaternary, a significant increase in mean body size (p-value = 0.003958), from 413 to 1036 mm, was detected for marine turtles (Figure 3c-d).

## 4. DISCUSSION

### 8.1. **Body size patterns and ancestral estimates: the effect of including fossils**

Despite previous controversies (e.g., Patterson, 1981), it has become increasingly clear that the paleontological record is crucial to answer macroevolutionary questions (Fritz et al., 2013; Louca & Pennell, 2020; Quental & Marshall, 2010). In particular, it is well-documented that the inclusion of fossils affects the estimation of ancestral states and evolutionary rates (e.g., Puttick, 2016; Slater et al., 2012). Nevertheless, extinct taxa is often neglected in such analyses, having so far been included in only one macroevolution study of turtle body size (although focused on a particular group, i.e., Testudinidae; Vlachos & Rabi, 2018).

As already noted by Jaffe et al. (2011), examining the evolution of body size in the fossil record of turtles might provide new insights not revealed by previous analyses. Based on a sample of 536 extinct taxa, our study was the first comprehensive attempt in that direction, confirming the impact of including fossils on estimates of both divergence times (Figure A1 and A2) and ancestral body sizes (Figure 1 and A3). The latter has been affected for most lineages assessed here, but to different degrees (Figure 1), with no directional effect on the reconstructions (Figure 1). This differs from the pattern seen in mammals, for which ancestors have considerably larger body sizes when estimated using fossils (e.g., Finarelli & Flynn, 2006; Finarelli & Goswami, 2013). For mammals, these results were explained by the widespread occurrence of directional evolution towards large body sizes from small ancestors (Alroy, 1999; Smith et al., 2010), a pattern not observed in turtles (see below).

### 4.2. Cope’s rule and directional trends of body size evolution

We found no evidence for directional patterns of body size evolution in turtles (Table 1), given that none of our trend-like models (either the uniform trend or the multi-trend models) received more support than the uniform OU model. This result is consistent with previous investigation by Moen (2006), which also found no support for Cope’s rule when analyzing extant cryptodires (even though directional evolution may be difficult to detect on extant-only datasets; Finarelli & Flynn, 2006; Schnitzler et al., 2017; Slater et al., 2012). Therefore, our results provide no support for the hypothesis that Cope’s rule explain the evolution of large body sizes in Testudinata. They also add to the growing evidence that directional body size evolution is rare among vertebrates (Benson et al., 2018; Godoy et al., 2019; Huttenlocker, 2014; Sookias et al., 2012), with the exception of mammals and pterosaurs (Alroy, 1998; Benson et al., 2014), once again challenging the generality of Cope’s rule.

### 4.3. The influence of environmental temperature on turtle body size evolution

The relationship between abiotic factors and body size has been extensively studied, with distinct vertebrate groups being differently affected by them, especially when it comes to comparing endothermic and ectothermic organisms (Angielczyk et al., 2015; Angilletta et al., 2004; Ashton & Feldman, 2003; Mousseau, 1997; Partridge & Coyne, 1997; van der Have & de Jong, 1996; Van Voorhies, 1996). Large-scale trends such as Bergmann’s rule (i.e., the tendency of having larger body sizes at higher latitudes within a species) may play an important role for within-species variation of body size in endotherms (Ashton et al., 2000; James, 1970; Zink & Remsen, 1986), and may explain patterns of maximum size during mammal evolution (Saarinen et al., 2014). Yet, results for ectothermic reptiles are less consistent (Angielczyk et al., 2015; Ashton & Feldman, 2003; Mousseau, 1997).

We evaluated the correlation between turtle body size distributions and paleotemperature variation through time. In general, no significant influence of temperature on mean, minimum or maximum body size in the group was found (Table A2). Similar results were reported for crocodylomorphs, for which no significant correlation between paleotemperature and body size (mean, maximum, and minimum values) was found when the entire group is analyzed, even though a strong association between both variables is observed when only the crown group is considered (Godoy et al., 2019). Therefore, although our results indicate no overall influence of paleotemperature on the through-time distribution of turtle mean, maximum, and minimum body sizes, we cannot rule out an influence of environmental temperatures at smaller temporal and phylogenetic scales. Indeed, environmental temperature has been a commonly proposed explanation for body size variation in different turtle clades and species, particularly affecting disparity, diversity, or distribution of less inclusive groups (e.g., Böhme, 2003; Ferreira et al., 2018; Georgalis & Kear, 2013; Vitek, 2012).

We did find a significant correlation between paleotemperature and turtle body size disparity (= standard deviation) during the Cenozoic, with periods of higher disparity associated with lower temperatures (or higher δ^18^O values; Figure 2). We suggest three potential explanations for this seemingly counterintuitive result. First, low temperatures might have restricted niche availability for turtles and, consequently, driven body size specialization – toward larger or smaller body sizes – to avoid competition, as seen in some extant lineages (Cunha et al., 2020; Pritchard, 2001). Conversely, colder and dryer environments could have increased availability of coastal habitats (by sea level drops), which has been associated with higher diversification rates in turtles (Thomson et al., 2021). Higher species richness might have also led to higher disparity levels in body size. Finally, the significant correlation might be an artefact from the coincidental drop in temperature over the Cenozoic and a continuous expansion in body size in turtles. Disparity constantly increases since the origin of the group, punctuated only by small drops (e.g., in the Oligocene and present time bins, Fig. 2). A slow, steady disparity increase has been also noted in cranial morphology by Foth & Joyce (2016), particularly during the Mesozoic in different lineages. That would also explain the stronger correlation with the Zachos et al. (2008) curve – restricted to Cenozoic δ^18^O values – in relation to that using Prokoph et al. (2001), which includes the period of increasing temperatures in the Mesozoic. In any case, it seems that environmental temperature did not play a major role in determining large-scale patterns of Testudinata body size variation through time, at least not when considering the entire group.

### 4.4. Body size evolution and ecological habitats

Our ANOVA results (Table 3) indicate a significant association between habitat preference and body size in turtles. This is seen in the phylogenetic ANOVA, specifically for freshwater and terrestrial habitats, indicating that evolutionary shifts of habitat correlate with directional evolutionary shifts of body size. Accordingly, through-time body size patterns for distinct habitat categories (Figure 3) can help understanding patterns observed in different turtle subgroups, which emphasizes the importance of independently examining each of the three main turtle ecologies: freshwater, terrestrial, and marine.

Since the Jurassic, most turtles have had freshwater ecologies (Figure 3b; Joyce & Gauthier, 2004), with these turtles keeping a fairly homogeneous body size disparity (= standard deviation) through time (Figure 3b). Their wide and constant disparity of body sizes might be explained by distinct evolutionary scenarios within such habitats (Jaffe et al., 2011), with different species, closely related or not, inhabiting several disparate freshwater environments (Bonin et al., 2006). For instance, different pleurodiran and cryptodiran lineages acquired resistance to estuarine or brackish water (Agha et al., 2018; Bower et al., 2016). Also, closely related taxa occupying the same areas are known to avoid competition through body size divergence, such as extant podocnemidids (Cunha et al., 2020) and trionychids (Pritchard, 2001).

Terrestrial and marine turtles, on the other hand, are represented by fewer lineages, with more restrict evolutionary histories. The earliest turtles were terrestrial, ranging from medium-to large-sized during the Mesozoic (Figure 3a-d). Meiolaniformes is the only of these stem-lineage groups to survive until recently (until the Holocene; Sterli, 2015), displaying large to gigantic body sizes, especially after the Mesozoic. Testudinids – the only extant lineage of terrestrial turtles – appeared in the fossil record during the Paleogene and remained relatively small until at least the end of that period (Figure 3c). The Eocene-Oligocene witnessed a peak of diversity, related to the origin of crown-Testudinidae (Lourenço et al., 2012; Vlachos & Rabi, 2018), after which the group spread from Eurasia to most of the world. Therefore, the recent high variation in body size within tortoises might be also related to the expansion of occupied habitats (as in freshwater turtles).

Extant marine turtles (Chelonioidea) exhibit low disparity, but remarkably large body sizes (Figure 3b), which might be related to morphological adaptations to a pelagic lifestyle, given that other groups of sea turtles (e.g., Thalassochelydia and Bothremydidae) are not as strongly associated with larger sizes and were probably not pelagic. The large size of marine turtles might also be explained by either physiological constraints (e.g., thermoregulation; Mrosovsky, 1980) or the need for higher dispersal abilities associated with migration (Jaffe et al., 2011). It has been previously proposed that thermoregulation and other physiological aspects (e.g., lung capacity while diving) play an important role in determining the larger body sizes of aquatic mammals and reptiles (Benson et al., 2012; Davis, 2014; Gearty et al., 2018; Gearty & Payne, 2020; Gutarra et al., 2022; Pyenson & Vermeij, 2016; Williams, 2001), by posing a minimum body size limit on these species. However, in the case of marine turtles, the fossil record shows that smaller species also existed in the past (Figure 3b and 4), with *Santanachelys gaffneyi* from the Early Cretaceous of Brazil as one of the oldest and smallest sea turtles (200 mm; Hirayama, 1998). Therefore, in the case of Testudinata, perhaps the lower body size limit imposed by physiological constraints was not as strict as those inferred for other secondarily aquatic tetrapods (e.g., mammals). On the other hand, the shell and the necessity to lay eggs on land possibly pose constraints on the maximum body sizes achieved by marine turtles (Benson et al., 2012), which are, in general, smaller than other Mesozoic marine reptiles and extant cetaceans (Benson et al., 2012; Smith & Lyons, 2011).

Finally, the shell might also constrain the maximum body size of turtles inhabiting terrestrial and semiaquatic environments (Golubović et al., 2017; Lyson et al., 2014). It could hamper turtles from attaining sizes as large as giant mammals and dinosaurs due to a different relation between weight and body size. Moreover, minimum body sizes in turtles are overall larger than that of the smallest lissamphibians, squamates, mammals, and birds. Endothermy could explain the smaller sizes of mammals and birds (Lovegrove, 2017), but not of lissamphibians or squamates. Hence, it is possible that the shell also imposes a lower body size limit to testudinatans.

## CONCLUSIONS

Turtle body sizes showed low disparity early in their evolutionary history. They reached substantial disparity only in the Early Cretaceous, concomitantly with the lowest mean body sizes. Habitat preference is only weakly linked to body size variation in turtles. Nevertheless, ecological transitions provide a partial explanation for differences in the body size distribution of turtle subgroups. Freshwater turtles show a constant range of body sizes and higher disparity through time, which might be related to the ecological diversity associated with these habitats. Body size in terrestrial turtles is explained by their ecological diversity, in addition to the higher dispersal ability in giant species. In sea turtles, upper and lower body size limits seem to be associated with physiological (e.g., thermoregulation) and morphological (e.g., the shell) constrains, as well as with adaptations to the pelagic lifestyle during the Quaternary.

We did not find support for a general trend-like process leading to larger body sizes, discarding Cope’s rule as an explanation for body size evolution in turtles. Also, we did not find a significant influence of paleotemperature on mean, maximum, and minimum body size. Although we found a significant, moderate correlation between temperature and body size disparity through time, this association might be an artefact caused by a join constant increasing of disparity and continuous drop in temperatures during the Cenozoic.

## DATA AVAILABILITY STATEMENT

The complete dataset, supertrees, and scripts used for the analyses in this study will be made available after peer review.

## ACKNOWLEDGEMENTS

This study was funded by São Paulo Research Foundation (FAPESP grants 2018/10276-7, 2019/06119-6, to BMF, 2022/05697-9, to PLG, and 2020/07997-4, toMCL), Swiss Government Excellence Scholarship (2021.0350 to BMF), and the National Science Foundation (NSF DEB 1754596 to PLG).

## APPENDIX

**Figure A1.**
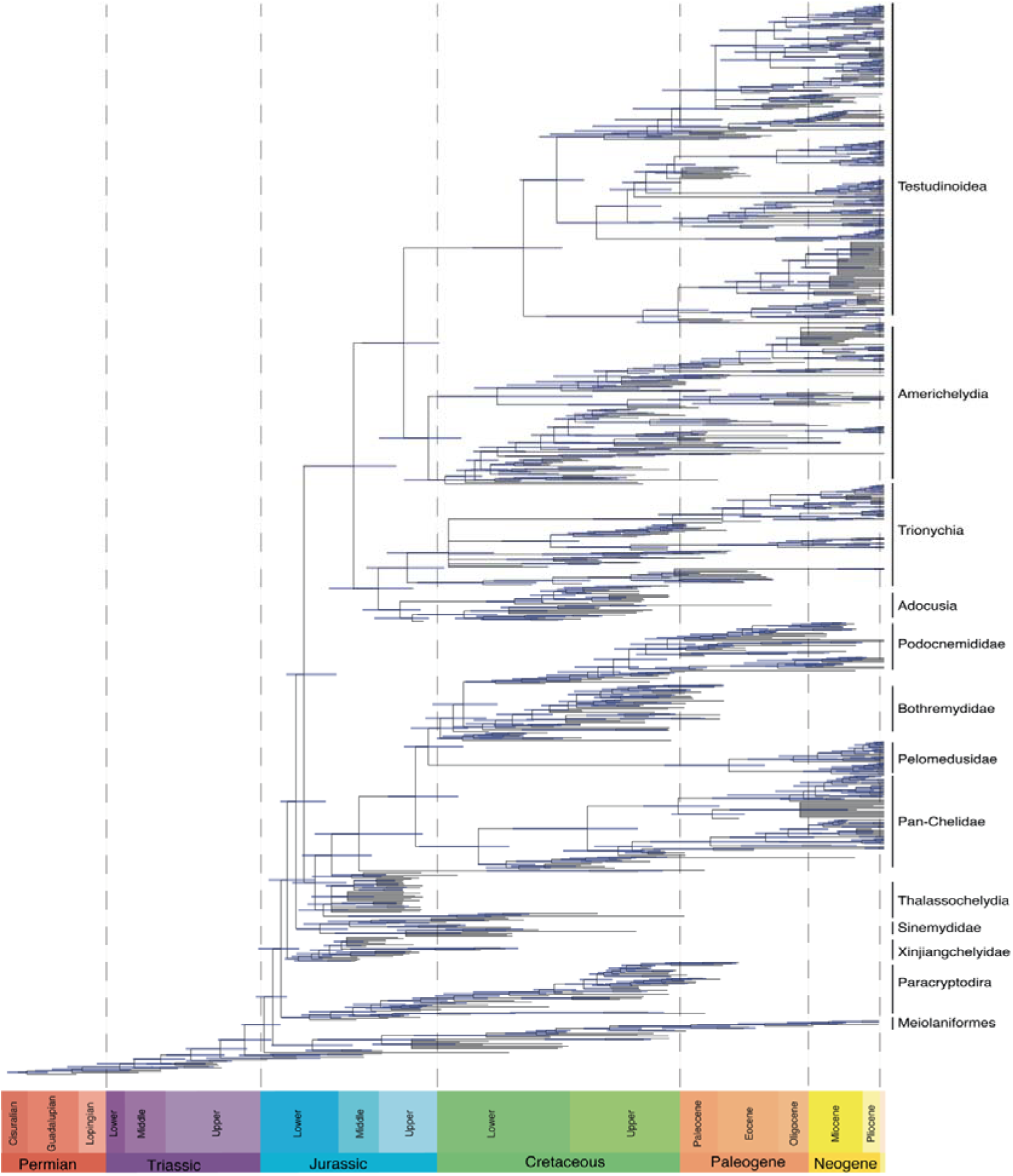
The **“**Ev19” time-calibrated supertree, represented by the 50% majority rule tree. Blue bars represent the 95% highest posterior density (HPD) age ranges for each node.

**Figure A2.**
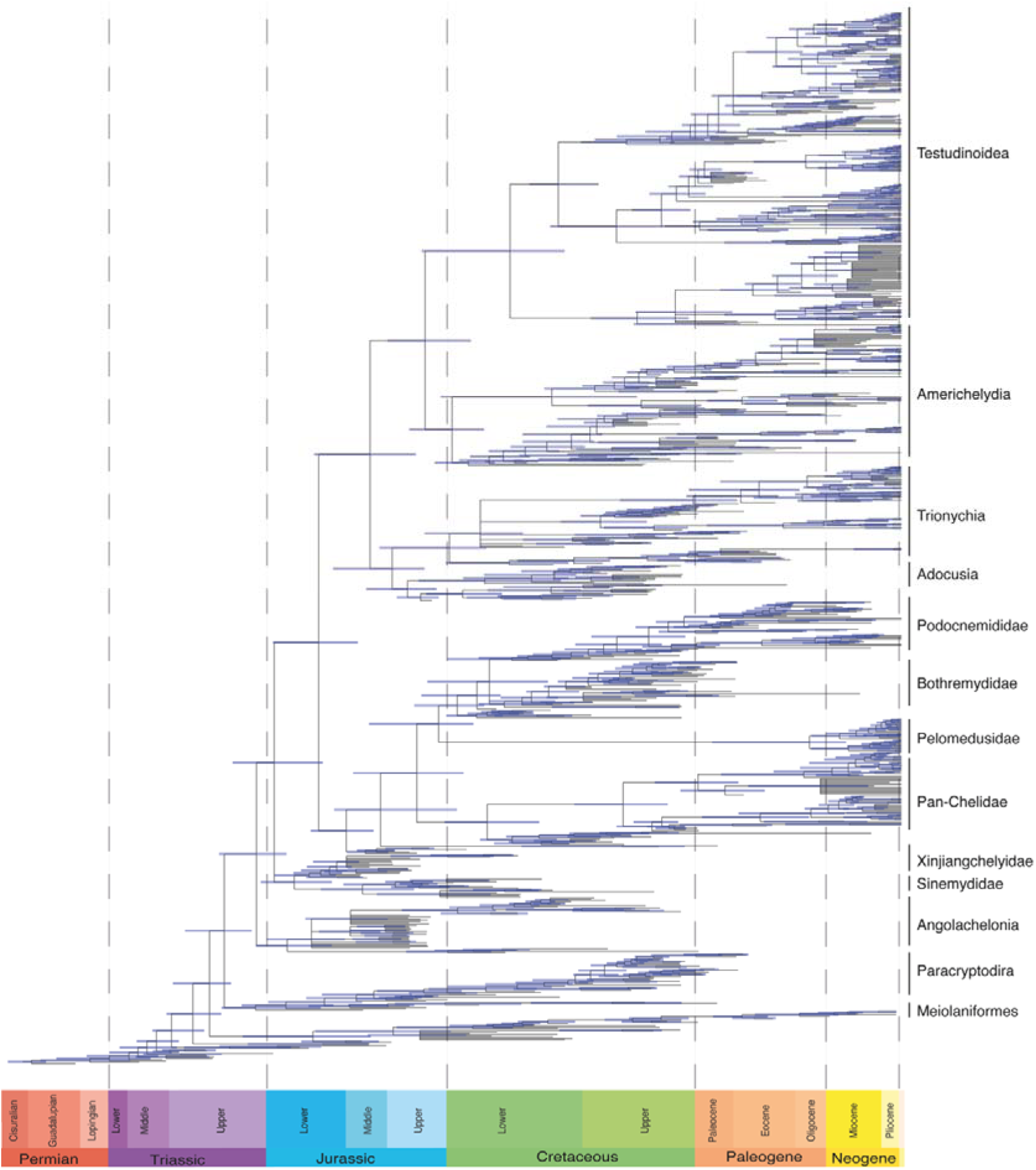
The “St18” time-calibrated supertree, represented by the 50% majority rule tree. Blue bars represent the 95% highest posterior density (HPD) age ranges for each node.

**Figure A3.**
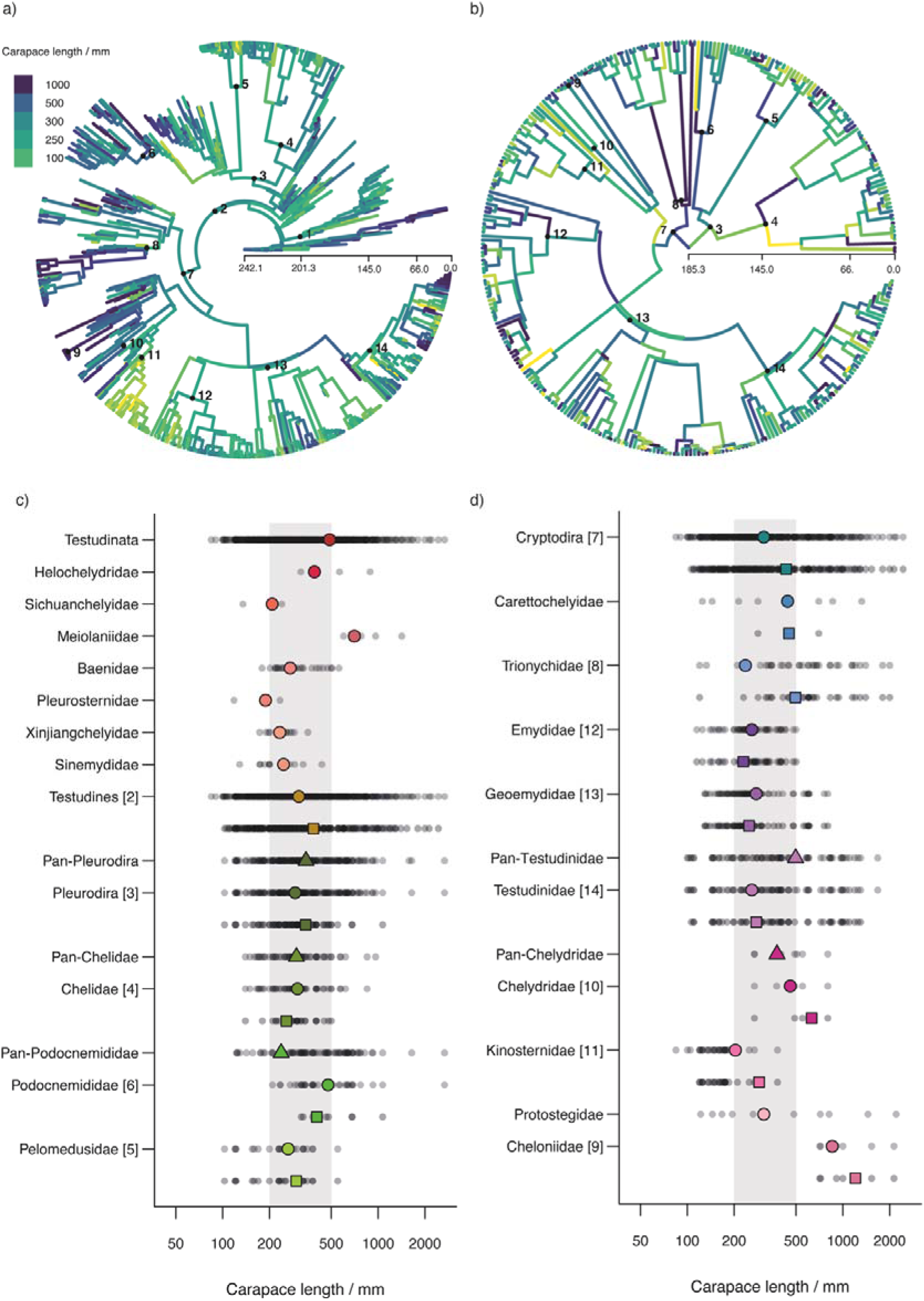
Ancestral body sizes (log_10_ maximum dorsal carapace length in millimeters) mapped onto Ev19 supertree, with (a) complete tree and (b) extant-only subtree. Ancestral body size for different taxonomic groups (c and d); small grey dots indicate all taxa within that lineage; colored triangles represent the ancestral estimates of stem-groups; circles represent ancestral estimates of the crown-groups and squares represent ancestral estimates of the crown-groups without the fossil taxa. Grey area indicates sizes between 200 and 500 mm. The numbers in the panels represent groups (same as in Figure 1).

**Table A1.** Sources of phylogenetic information for supertrees construction.

| Group | References |
| --- | --- |
| Testudinata | Evers et al. (2019); Sterli et al. (2018).<br>*Joyce (2017): used to include taxa that were not sampled by the above topologies. |
| Meiolaniformes | Sterli et al. (2015); Sterli et al. (2018).<br>*In “Ev19”: the position of *Chubutemys* and *Kallokibotion* were defined according to Evers et al. (2019). |
| Helochelydridae | Joyce (2017)<br>* *Aragochersis*: according to Pérez-García et al. (2019). |
| Sichuanchelyidae | Joyce (2017) |
| Compsemydidae | Joyce & Anquetin (2019)<br>* *Peltochelys duchastelii*: according to Joyce & Rollot (2020). |
| Paracryptodira | Joyce & Rollot (2020)<br>*position of *Dinochelys*, *Compsemydidae*, *Uluops* e *Baenoidea*. |
| Baenidae | Joyce & Lyson (2015)<br>*Eubaeninae (Lyson et al., 2019) + “*Baena*” *escavada* (Joyce & Lyson, 2015). |
| Pleurosternidae | Joyce & Rollot (2020)<br>*Siamochelys peninsularis*: according to Sterli et al. (2018).<br>*sister group of Paracryptodira |
| Xinjiangchelyidae | Pérez-García et al. (2014) *Xinjiangchelys (less inclusive clade) according to Evers et al. (2019).<br>Sampled made by Evers & Benson (2019) and Evers et al. (2019) include: *Xinjiangchelys radiplicatoides*, *Xinjiangchelys wusu*, *Annemys* IVPP V18106, *Annemys levensis*, and *Annemys latiensi*. *A. latiensi* and *A. levensis* are included in Pérez-García et al. (2014) as *Xinjiangchelys latiensi* e *Xinjiangchelys levensis*, but other *Xinjiangchelys* in Evers et al. (2019) are not. Because *Xinjiangchelys* is monophyletic in Pérez-García et al. (2014) (if *Larachelus* and *Brodiechelys* are included), we assume the Evers et al. (2019)’s topology includes only taxa from this clade (i.e., clade that includes *Xinjiangchelys*, *Larachelus* e *Brodiechelys*, but not *Protoxinjiangchelys* or *Tienfuchelys*), thus the relationship between other xinjiangchelyidids follow Pérez-García et al. (2014). |
| Sinemydidae | Shao et al. (2018); Evers & Benson (2019).<br>* *Hoyasemys jimenezi*: according to Pérez-García et al. (2012).<br>* *Galvechelone lopezmartinezae*: according to Pérez-Garcia & Murelaga (2012). |
| Angolachelonia | Evers et al. (2019) |
| Thelassochelydia | Anquetin et al. (2017)<br>* In “St18” *Santanachelys* is included within Eurysternidae. |
| Sandownidae | Evers et al. (2019) |
| Platycheilyidae | López-Conde et al. (2017) |
| Dortokidae | Cadena & Joyce (2015) |
| Pan-Chelidae | Holley et al. (2020); Pereira et al. (2017).<br>* *Linderochelys* and *Salamanchelys*: according to Hermanson et al. (2020).<br>* *Yaminuechelys sulcipeculiaris*: according to Orizabala et al. (2020). |
| Pelomedusidae | Pereira et al. (2017) ; Petzold, et al. (2014). |
| Pan-Podocnemididae | Hermanson et al. (2020) |
| Adocidae | Syromyatnikova & Danilov (2013)<br>* *Adocus inexpectatus*: according to Danilov, et al. (2013). |
| Nanhsiungchelyidae | Tong & Li (2019).<br>* *Basilemys gaffney*, *Basilemys morriensis* and *Zangerlia testudinomorpha*: Mallon & Brinkman (2018). |
| Pan-Carettochelys | Havlik et al. (2014). |
| Trionychia | Pereira et al. (2017), Brinkman et al. (2017; implied weighting tree); other added according to: Vitek & Joyce (2015) and Georgalis & Joyce (2017).<br>* Plastomenidae according to Joyce, Lyson, & Sertich (2018).<br>* *Axestemys* according to Vitek & Joyce (2015).<br>* *Apalone amorensis*: according to Valdes et al. (2017).<br>* *Aspideretoides foveatus* and *Gobiapalone orlovi*: according to Brinkman et al. (2017). |
| Pan-Testudinoidea | Vlachos (2018); Pereira et al. (2017) (general topology).<br>**Pan-Emydidae:** Vlachos (2018); Pereira et al. (2017).<br>*Polytomy within *Pseudemys*, *Graptemys* and *Trachemys*<br>**Pan-Geoemydidae:** Vlachos (2018); Pereira et al. (2017).<br>* *Banhxeochelys* and *Guangdongemys* according to Garbin et al. (2019).<br>* *Hardella siamensis*: according to Claude et al. (2007). |
|  | \**Pangshura tatrotia*: according to Joyce & Lyson (2010).<br>**Pan-Testudinidae**: Vlachos & Rabi (2018), Vlachos (2018), and Pereira et al. (2017).<br>*Extant taxa were included following Pereira et al. (2017); extinct taxa were positioned according to Vlachos & Rabi (2018) and Vlachos (2018).<br>* *Gopherus*: Vlachos (2018).<br>* Polytomy within *Chelonoidis*<br>**Geochelona**: Pereira et al. (2017), Vlachos & Rabi (2018), Pérez-García et al. (2020).<br>**Pan-Testudona**: Vlachos & Tsoukala (2016)<br>\**Impregnochelys* and *Gigantochersina*: according to Pérez-García et al. (2020). |
| Pan-Chelydridae | Joyce (2016) |
| Pan-Kinosternoidea | Joyce & Bourque (2016)<br>\**Kinosternon*: polytomy (we added taxa from Pereira et al. (2017)’s topology).<br>\**Yelmochelys rosarioae*: according to Brinkman, Aguillon-Martinez et al. (2016).<br>\**Lutemys warren*: according to Lyson et al. (2017).<br>*Kinosterninae: Joyce & Bourque (2016). |
| Pan-Chelonioidea | Gentry et al. (2019), and Evers et al. (2019).<br>\**Mexichelys*, *Argillochelys antiqua* and *Procolpochelys*: according to Zvonok & Danilov (2017).<br>\**Prionochelys*, *Euclastes wielandi*\*\*, *Asmodochelys*: according to Gentry et al. (2019).<br>\**Osonachelus decorata*: according to Lapparent de Broin et al. (2014).<br>\**Allopleuron qazaqstanense* and *Allopleuron lipsiensis* according to Karl et al. (2012).<br>\**Rhinochelys amaberti* according to Scavezzoni & Fischer (2018).<br>*Ctenochelyidae: Evers et al. (2019).<br>\**Santanachelys*: Protostegidae, according to Gentry et al. (2019) Evers et al. (2019), and Scavezzoni & Fisher (2018), closely related to *Solnhofia* (Thalassochelyidae according to Anquetin et al. 2017 and Evers et al. 2019), Protostegidae become more basal, within Thalassochelyidae.<br>*The position of Cheloniidae and Ctenochelyidae according to Evers et al. (2019).<br>\**Carolinochelys* and *Trachyaspis* closely related to Cheloniidae is supported by Weems & Brown (2014) and Zvonok & Danilov (2017).<br>\**Pacifichelys*, *Erquelinnesia*, *Tasbacka spp.*, “*Argillochelys*” *africana* and *Euclastes* is supported partially by Weems & Brown (2014) and based on Zvonok & Danilov (2017).<br>*Lophochelyinae as sister group of those groups, excluding *Toxochelys*, and *Mexichelys* is supported by Parham & Pyenson (2010), Weems & Brown (2014) and Zvonok & Danilov (2017). |

**Table A2.** Results of generalized least squares (tsGLS) and ordinary least squares (OLS) regressions using turtle mean, maximum, and minimum body size and paleotemperature data. Body size (log_10_ maximum dorsal carapace length in millimeters) and paleotemperature (δ^18^O data as a proxy for paleotemperature from two different sources: Prokoph et al. [2008] and Zachos et al. [2008]) data was divided into time bins (24 time-bins when the Prokoph et al. [2008] δ^18^O data was used and 10 time-bins when the Zachos et al. [2008] data was used). Abbreviation: **sd.**, standard deviation; **p.R^2^**, Nagelkerke pseudo R-squared; **Var.**, variable; **interc.**, intercept; **Coef.**, coefficient.

|  | tsGLS |  |  |  |  |  |  | OLS |  |  |  |  |  |
| --- | --- | --- | --- | --- | --- | --- | --- | --- | --- | --- | --- | --- | --- |
|  | Phi | p.R<sup>2</sup> | AIC | Var. | Coef. | t-value | p-value | p.R<sup>2</sup> | AIC | Var. | Coef. | t-value | p-value |
| Body size<br>mean<br>~ $\delta^{18}\text{O}$<br>(Prokoph) | 0.761 | -0.004 | -57.426 | interc. | 2.494 | 46.05 | <0.000 | -0.009 | -40.37 | interc. | 2.47 | 69.942 | <0.000 |
| | | | | $\delta^{18}\text{O}$ | -0.01 | -1.140 | 0.266 | | | $\delta^{18}\text{O}$ | -0.025 | -1.061 | 0.300 |
| Body size<br>mean<br>~ $\delta^{18}\text{O}$<br>(Zachos) | 0.121 | -0.0003 | -36.809 | interc. | 2.558 | 132.713 | <0.000 | -0.0004 | -38.663 | interc. | 2.558 | 144.934 | <0.000 |
| | | | | $\delta^{18}\text{O}$ | -0.004 | -0.452 | 0.663 | | | $\delta^{18}\text{O}$ | 0.01 | -0.508 | 0.626 |
| Body size<br>max.<br>~ $\delta^{18}\text{O}$<br>(Prokoph) | 0.859 | -0.038 | -14.857 | interc. | 3.12 | 16.439 | <0.000 | 0.185 | 13.608 | interc. | 3.132 | 28.813 | <0.000 |
| | | | | $\delta^{18}\text{O}$ | 0.046 | 1.130 | 0.271 | | | $\delta^{18}\text{O}$ | 0.085 | 1.166 | 0.256 |
| Body size<br>max.<br>~ $\delta^{18}\text{O}$<br>(Zachos) | 0.469 | -0.043 | -8.991 | interc. | 3.163 | 29.724 | <0.000 | -0.161 | -8.652 | interc. | 3.130 | 39.545 | <0.000 |
| | | | | $\delta^{18}\text{O}$ | 0.070 | 1.348 | 0.214 | | | $\delta^{18}\text{O}$ | 0.085 | 1.923 | 0.091 |
| Body size<br>min.<br>~ $\delta^{18}\text{O}$<br>(Prokoph) | 0.937 | -0.011 | -45.143 | interc. | 2.155 | 12.262 | <0.000 | -0.237 | -17.639 | interc. | 1.978 | 34.891 | <0.000 |
| | | | | $\delta^{18}\text{O}$ | 0.03 | -1.417 | 0.171 | | | $\delta^{18}\text{O}$ | | -2.661 | 0.014 |
| Body size<br>min.<br>~ $\delta^{18}\text{O}$<br>(Zachos) | 0.420 | -0.0003 | -26.379 | interc. | 1.988 | 46.845 | <0.000 | -0.00001 | -27.545 | interc. | 1.994 | 64.800 | <0.000 |
| | | | | $\delta^{18}\text{O}$ | 0.007 | 0.33 | 0.75 | | | $\delta^{18}\text{O}$ | -0.001 | -0.048 | 0.963 |

